# Efficient inactivation of SARS-CoV-2 by WHO-recommended hand rub formulations and alcohols

**DOI:** 10.1101/2020.03.10.986711

**Authors:** Annika Kratzel, Daniel Todt, Philip V’kovski, Silvio Steiner, Mitra L. Gultom, Tran Thi Nhu Thao, Nadine Ebert, Melle Holwerda, Jörg Steinmann, Daniela Niemeyer, Ronald Dijkman, Günter Kampf, Christian Drosten, Eike Steinmann, Volker Thiel, Stephanie Pfaender

## Abstract

The recent emergence of Severe acute respiratory syndrome coronavirus 2 (SARS-CoV-2) causing COVID-19 is a major burden for health care systems worldwide. It is important to address if the current infection control instructions based on active ingredients are sufficient. We therefore determined the virucidal activity of two alcohol-based hand rub solutions for hand disinfection recommended by the World Health Organization (WHO), as well as commercially available alcohols. Efficient SARS-CoV-2 inactivation was demonstrated for all tested alcohol-based disinfectants. These findings show the successful inactivation of SARS-CoV-2 for the first time and provide confidence in its use for the control of COVID-19.

**Importance:** The current COVID-19 outbreak puts a huge burden on the world’s health care systems. Without effective therapeutics or vaccines being available, effective hygiene measure are of utmost importance to prevent viral spreading. It is therefore crucial to evaluate current infection control strategies against SARS-CoV-2. We show the inactivation of the novel coronavirus for the first time and endorse the importance of disinfectant-based hand hygiene to reduce SARS-CoV-2 transmission.

## Introduction

SARS-CoV-2 is the third highly pathogenic human CoV to have crossed the species barrier into humans during the last 20 years^1,2,3^. SARS-CoV-2 infection is associated with coronavirus disease 2019 (COVID-19) which is characterized by severe respiratory distress, fever and cough, leading to a high percentage of fatalities, especially in the elderly or patients with comorbidities^4^. As of March 6^th^ 2020 there are 101604 globally confirmed cases and related 3460 deaths^5^. Health care experts suspect that a global pandemic is inevitable mainly because human-to-human transmission of SARS-CoV-2 is very efficient and infected individuals can transmit the virus without or with only mild symptoms^4^. Given that so far, no therapeutics or vaccines are available, virus containment and prevention of infection are of highest priority. Effective hand hygiene is crucial to limit virus spread. Therefore, easily available but efficient disinfectants are crucial. The World Health Organization’s ‘Guidelines for Hand Hygiene in Health Care’ suggests two alcohol-based formulations for hand sanitization to reduce pathogen infectivity and spreading. These recommendations are based on fast-acting and broad-spectrum of microbicidal activity, as well as easy accessibility and safety^3^. We have previously shown that WHO formulation I and II were able to inactivate the closely related SARS-CoV and MERS-CoV^6^. So far, recommendations to inactivate SARS-CoV-2 were only translated from findings with other coronaviruses^7^. To evaluate if alcohol-based disinfectants are also efficient for the inactivation of SARS-CoV-2, we tested different concentrations of WHO formulation I and II, as well as the alcohols ethanol and 2-propanol for their virucidal activity.

## Results

SARS-CoV-2 was highly susceptible to the WHO formulations (**Fig. 1**). WHO formulation I, based on 85 % ethanol, efficiently inactivated the virus with reduction factors (RFs) of ≤ 5.9 and concentrations between 40 % – 80 % (**Fig. 1A**). Subsequent regression analysis revealed similar inactivation profiles compared to SARS-CoV, MERS-CoV and bovine CoV (BCoV), which is often used as surrogate for highly pathogenic human CoVs (**Fig. 1A**). WHO formulation II, which is based on 75 % isopropanol, demonstrated a better virucidal effect at low concentrations, with complete viral inactivation and RFs of ≤ 5 at a minimal concentration of 30 % (**Fig. 1B**). The regression analysis showed an inactivation profile of SARS-CoV-2, which was in between SARS-CoV, BCoV and MERS-CoV **(Fig. 1B)**.

**Figure 1.**
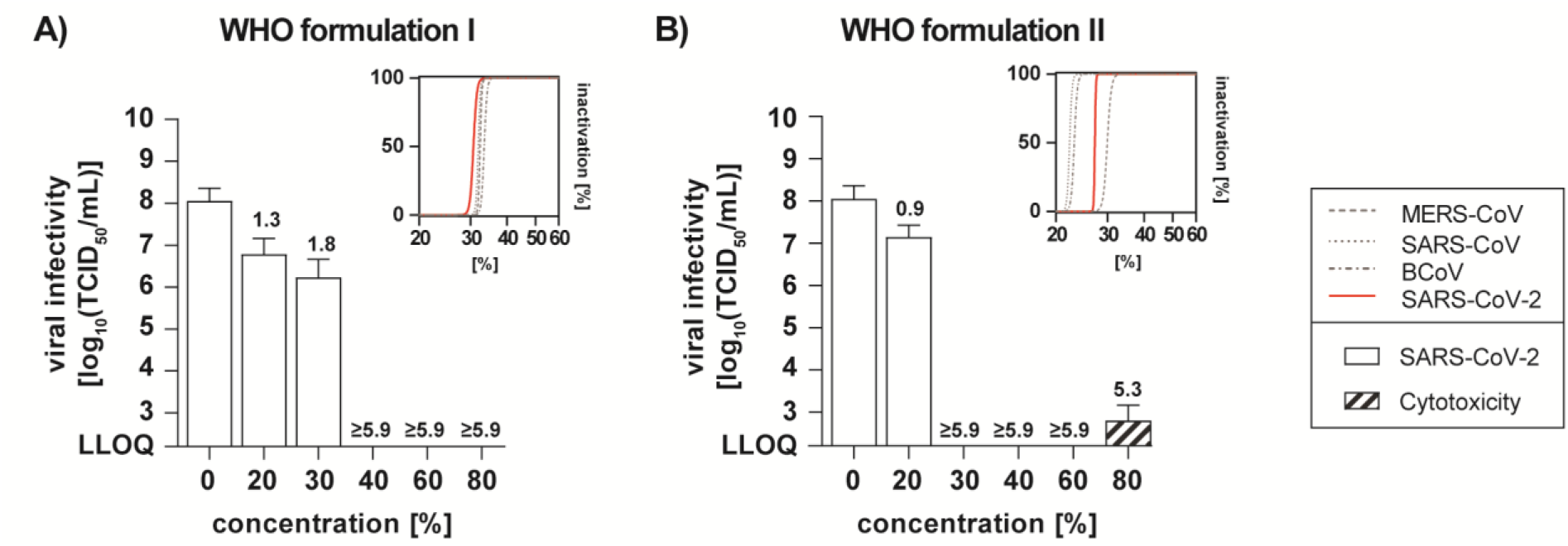
Virucidal activity of WHO formulations I (85 % ethanol) and II (75 % 2-propanol) against SARS-CoV-2. WHO formulations I (**A**) and II (**B**) were tested for their efficacy in inactivating SARS-CoV-2. The concentrations of the WHO formulations ranged from 0 % to 80 % with an exposure time of 30 seconds. Viral titers are displayed as 50 % tissue culture infectious dose 50 (TCID50/mL) values. Cytotoxic effects are displayed as dashed bars are and were calculated analogous to virus infectivity. RFs are included above the bar. The mean of two - three independent experiments with standard deviation are shown. LLOQ: lower limit of quantification. Top inserts: Regression analysis of the inactivation of SARS-CoV-2, bovine CoV (BCoV), SARS-CoV and MERS-CoV by WHO formulation I (**A**) and II (**B**). Depicted is the percentage of inactivation at different concentrations.

Next, we addressed the susceptibility of SARS-CoV-2 against the individual components of the WHO recommended formulations which are also the main ingredients of commercially available hand disinfections. Both alcohols, ethanol (**Fig. 2A**) and 2-propanol (**Fig. 2B**) were able to reduce viral titers in 30 s exposure to background levels with RFs between ≤ 4.8 and 5.9 after 30 sec. Furthermore, we could show that a minimal concentration of 30 % ethanol or 2-propanol is sufficient for viral inactivation (**Fig. 2**).

**Figure 2.**
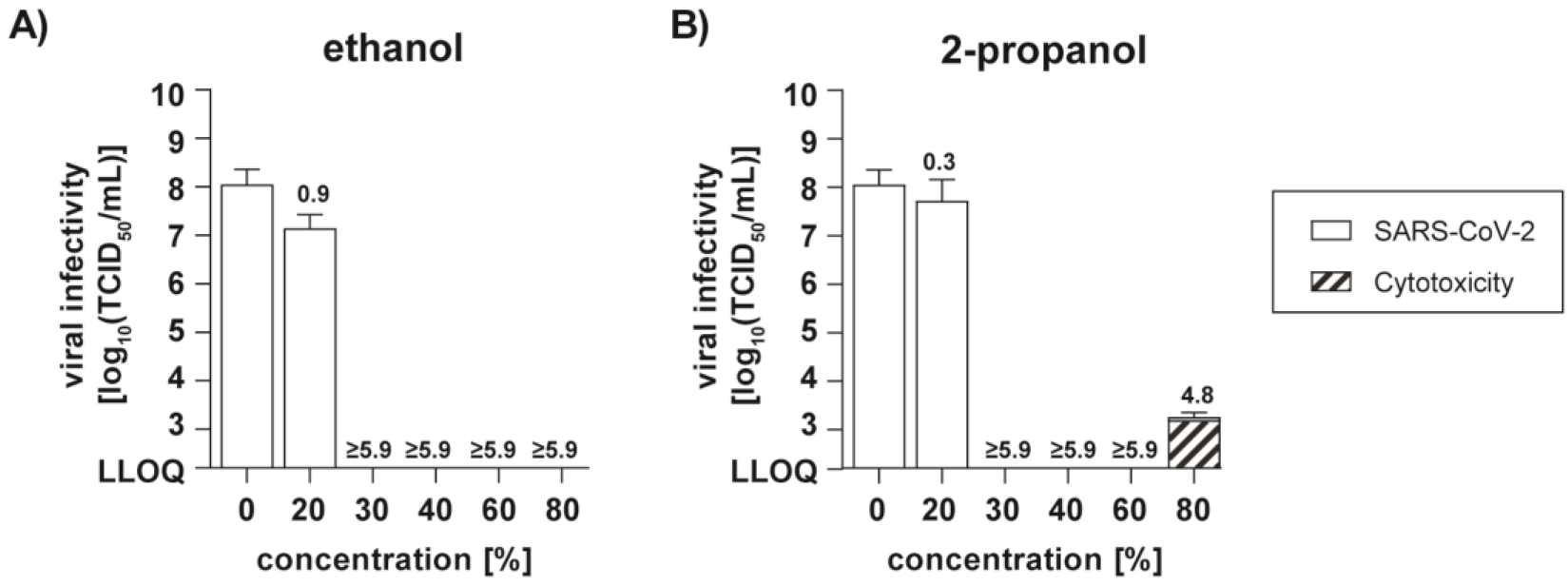
Effect of alcohols on SARS-CoV-2. Commercially available ethanol (**A**), or 2-propanol (**B**) were tested for their efficacy in inactivating SARS-CoV-2. The biocide concentrations ranged from 0 % to 80 % with an exposure time of 30 seconds. Viral titers are displayed as 50 % tissue culture infectious dose 50 (TCID_50_/mL) values. Cytotoxic effects are displayed as dashed bars are and were calculated analogous to virus infectivity. LLOQ: lower limit of quantification. RFs are included above the bar. Dashed line: limit of detection. The mean of two - three independent experiments with standard deviation are shown.

## Discussions

This study shows that SARS-CoV-2 can be efficiently inactivated by both WHO formulations implicating their use in health care systems and viral outbreak situations. Notably, both tested alcohols, ethanol and 2-propanol were efficient in inactivating the virus in 30 s at a minimal final concentration of at least 30 %. Alcohol constitutes the basis for many hand rubs routinely used in health care settings. Our findings are therefore of utmost importance in the current outbreak situation to minimize viral transmission and maximize virus inactivation.

## Material and Methods

### Viral strains and cell culture

SARS-CoV-2 (SARS-CoV-2/München-1.1/2020/929) stocks were propagated on VeroE6 cells (kindly provided by M-Müller/ C. Drosten, Charité, Berlin, Germany). VeroE6 cells were cultured in Dulbecco’s modified minimal essential medium (Gibco) supplemented with 10 % heat inactivated fetal bovine serum (Gibco), 1 % non-essential amino acids (Gibco), 100 μg/mL Streptomycin and 100 IU/mL Penicillin (Gibco) and 15 mM HEPES (Gibco).

### Chemicals

WHO I formulation consists of 85 % ethanol (v/v), 0.725 % glycerol (v/v) and 0.125 % hydrogen peroxide (v/v). The isopropyl-based formulation, WHO II, contains 75 % isopropanol (w/w), 0.725 % glycerol (v/v) and 0.125 % hydrogen peroxide (v/v)^8^. In addition, ethanol (CAS 64-17-5), and 2-propanol (CAS 67-63-0) were investigated.

### Quantitative Suspension Test and Virus Titration

Virucidal activity studies were performed with a quantitative suspension test with 30 seconds exposure time^3^. Briefly, one part virus suspension was mixed with one part organic load (0.3 % bovine serum albumin [BSA] as interfering substance) and eight parts disinfection solution of different concentrations. Following 30 seconds exposure, samples were serially diluted and the TCID50/mL values were determined by crystal violet staining and subsequent scoring the amounts of wells displaying cytopathic effects. TCID_50_ is calculated by the Spearman & Kärber algorithm as described^9^. Cytotoxic effects of disinfectants were monitored by crystal violet staining and optical analysis for altered density and morphology of the cellular monolayer in the absence of virus and were quantified analogous to the TCID_50_/mL of the virus infectivity.

### Statistical Analysis

Dose-response curves (normalized virus inactivation [%] *vs*. log (disinfectant concentration [%]) were determined using nonlinear regression using the robust fitting method on the normalized 50 % tissue culture infectious dose (TCID_50_) data implemented in GraphPad Prism version 8.0.3 for Windows. Reference curves for SARS-CoV, MERS-CoV and BCoV were plotted based on previously published data^6^. The mean TCID50 and standard deviations of means were assessed from 2-3 individual experiments. Outlier were identified using Grubb’s test (GraphPad Prism). Reduction factors (RF) for each treatment condition were calculated as follows:

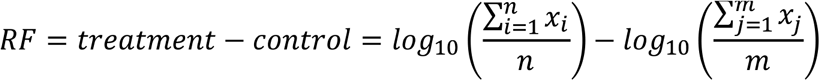

## Acknowledgments

We would like to thank all members of Institute of Virology and Immunology, Bern and the Department for Molecular & Medical Virology for helpful suggestions and discussions.

